# Data-driven oscillatory network modeling with condition-dependent coupling laws: Identifying directed neural interactions in working memory–attention dynamics

**DOI:** 10.64898/2026.07.06.736523

**Authors:** Mika Ohkawa, Ying Joey Zhou, Saskia Haegens, Matin Jafarian

## Abstract

Learning new information in the presence of distracters and changing conditions requires the ability to adapt. In the brain, this adaptive capability has been linked to dynamic interactions between attention and working memory, which enable the selective filtering of irrelevant input while preserving behaviorally relevant information. Specific neural oscillations have been implicated in this process.

Here, we introduce a phenomenological data-driven framework for oscillatory network modeling that learns condition-dependent coupling laws directly from neural recordings and enables inference of condition-dependent directed pathways. We apply our approach to magnetoencephalography (MEG) data collected while participants performed a working-memory task with and without distracters. Recall dynamics in the non-distracter condition are first modeled using a linear oscillatory network in which each region of interest is represented by two alpha-band harmonic oscillators. We use universal differential equations (UDE), an extension of neural differential equations, to capture distracter-induced changes in coupling laws. Symbolic regression is then used to interpret the modifications identified by UDE as nonlinear functions, and an additional method is proposed to identify the directed pathway from the newly emerging nonlinear terms in the dynamics of brain regions of interest.

Despite inter-subject variability, working memory recall data from all four participants examined under distraction showed the emergence of a pathway from the dorsolateral prefrontal cortex (dlPFC) to the primary visual cortex (V1). This finding is consistent with the established role of the dlPFC in cognitive control and suggests that distracter processing recruits a directed interaction from prefrontal to visual regions. More broadly, our results illustrate that combining linear models whose parameters are learned from the data with universal differential equations augmented by interpretability methods enables the identification of condition-dependent coupling laws, their representation as interpretable mathematical functions, and the discovery of candidate directed pathways underlying adaptive changes in oscillatory networks without requiring strong prior assumptions about the underlying mechanisms.

## 1 Introduction

The ability to learn and maintain performance despite external distractions and changing sensory inputs requires a flexible, adaptive system. In the brain, this adaptive capability has been linked to dynamic interactions between attention and working memory, which enable the selective filtering of irrelevant input while preserving behaviorally relevant information. These processes have been extensively studied in psychology and cognitive science, and more recently in neuroscience [1, 2]. Many biological dynamical systems exhibit oscillatory activity whose underlying generative dynamics change across varying conditions. Indeed, neuroimaging studies [3, 4] have shown the underlying role of brain oscillations in working memory and attention, especially by modulating the interplay between these cognitive processes. Although available data have increased with advances in measurement and imaging technology, biophysical mechanisms remain largely unclear [5]. Our goal is to develop a phenomenological, data-driven framework for oscillatory network modeling that captures condition-dependent mechanisms directly from data and enables the inference of directed pathways that emerge across conditions. A central motivation for this work is to understand the mechanisms governing interactions between visual attention and working memory.

Attention has long been seen as a mechanism for allocating the brain’s limited sensory-processing resources [2]. Early theories of attention explained it in terms of changes in neuronal firing rates, leading to rate-based models such as the biased-competition principle, in which competing visual signals are biased toward behaviorally relevant stimuli [6, 7]. Building on these ideas, cognitive models of working memory (WM) identify attention as a key mechanism for filtering sensory information and maintaining task-relevant memory contents [8, 9]. Advancing this perspective, more recent neuroimaging studies suggest that oscillatory neuronal activity underlies these mechanisms. In particular, alpha-band modulation has been linked to WM recall performance [10] and distracter suppression, with increased alpha activity observed in regions processing task-irrelevant stimuli [3, 4]. Although findings remain partly inconsistent [11, 12], current theories suggest that alpha synchronization may reduce neuronal excitability and suppress competing signals [13,14]. Together, these findings suggest that low-frequency oscillations support top-down control, with alpha activity regulating the enhancement or suppression of sensory information in working memory.

But how do relevant brain regions involved in the interplay of WM and attention dynamically interact? This question has motivated the development of generative dynamic models, including oscillatory network models, ranging from detailed biophysical neuronal networks to abstract phenomenological oscillator models [15, 16]. Existing white-box models, such as biologically plausible neuronal networks and neural mass models, are typically hypothesis-driven [17–19]. They often include many parameters, which can be computationally expensive, difficult to fit to the data, and limit data-driven exploration and the interpretation of the underlying dynamics [20]. Methods such as dynamic causal modeling [21] partly address these issues by combining biologically informed structures with Bayesian parameter estimation, though they also require a large model bank and remain primarily suited to hypothesis testing rather than exploratory modeling [22]. Moreover, these methods mainly explain changes in connectivity, expressed by positive or negative scalar, between brain regions which can be difficult to interpret when the experiment aims to compare mechanisms across conditions, such as information encoding under conditions with versus without distracters. This motivates the use of function identification approaches.

Recent advances in machine learning, particularly scientific machine learning, leverage neural networks as universal function approximators [23]. Scientific machine learning enables the incorporation of prior scientific knowledge to constrain learning and improve interpretability [24, 25]. A key development in this area is the Neural Ordinary Differential Equation (NODE) framework [26], in which system dynamics are represented by a neural network embedded in a differential equation. Universal Differential Equations (UDEs) further extend this approach by decomposing the system dynamics into a sum of known and unknown parts, and using neural networks only to model the unknown part of the dynamics [24]. This framework reduces model complexity while retaining flexibility, making it well-suited to exploratory dynamical modeling in neuroscience. UDEs have been successfully applied across a wide range of scientific domains. In physics, they have recovered known black hole dynamics and uncovered new equations of motion from gravitational wave data [27]. In electrochemical engineering, they outperform traditional models of lithium-ion battery degradation [28], while in epidemiology, they have been used to identify the effects of quarantine policies on COVID-19 transmission dynamics [29]. To the best of our knowledge, however, the application of UDEs for neuronal dynamics remains largely unexplored, despite their potential to uncover novel neural mechanisms. While UDEs provide a powerful tool for capturing nonlinear dynamics, their mathematical interpretation should be carried out using techniques such as symbolic regression. These interpretations yield complex mathematical expressions, such as sums of monomials, that may not be immediately insightful regarding dynamical properties. Consequently, improving the interpretability of UDEs may require combining them with novel interpretive methods.

We present a phenomenological, data-driven oscillatory network modeling framework to capture condition-dependent mechanisms directly from data. We use data from a previous working memory-visual attention experiment measured with magnetoencephalography (MEG). Our modeling is partially informed by established neuroscience knowledge, such as selection of regions of interest and frequency bands, while retaining sufficient flexibility to uncover novel mechanisms. We implement our algorithm to capture recall dynamics corresponding to brain data from four participants who performed a working-memory task in conditions with and without distracters [3]. To do this, we first propose a linear oscillatory network. We allocate two harmonic oscillators to each region of interest in the alpha band, to model WM recall dynamics and identify model parameters for the condition without distracters. We then use UDEs to capture functional coupling changes for the same participants when distracters were presented [3]. Thereafter, we use symbolic regression to identify the changes in couplings in terms of nonlinear functions and propose a method, based on assessing which region is causing new nonlinear terms to the dynamics of their coupled regions, to identify the candidates for emerging attention-dependent pathway. We find that the pathway from the dorsolateral prefrontal cortex (dlPFC) to the primary visual cortex (V1) is activated in the condition with distracters, consistent with prior work on the role of dlPFC in cognitive control. We conclude by discussing the novelty of our approach, its limitations as well as potential applications to broader questions in neuroscience.

## 2 Methods

We develop a phenomenological data-driven oscillatory network modeling algorithm that learns condition-dependent coupling laws among brain regions of interest directly from neural recordings. A central motivation for this work is to understand the mechanisms governing interactions between visual attention and working memory. This section presents the problem statement, our proposed approach and preliminaries required for modeling.

### 2.1 Data and experimental paradigm

The data used in this project were taken from a human MEG study [3] focused on how alpha-band activity in sensory cortex relates to distracter suppression during WM tasks. The participant’s task was to memorize a visually presented array of six digits, and report the digit at the probed spatial location in each trial. Each trial consisted of three phases: encoding, maintenance and recall (Figure 1). At the start of the trial, a visual cue was provided. In this work, we focus on two cues indicating either the presence or absence of the distracter. This phase was followed by the encoding phase, during which the target array was presented. During the maintenance phase, distracters appeared (or not) dependent on the distracter condition. During the recall phase, the participant was probed to recall the target in exactly the same manner across the two conditions. Thus, the observed change in the alpha power spectra [3] between the recall in these two conditions can be ascribed to an internal change in dynamics triggered by distracter presence. This internal change is assumed to be the activation of attention mechanisms.

**Fig 1.**
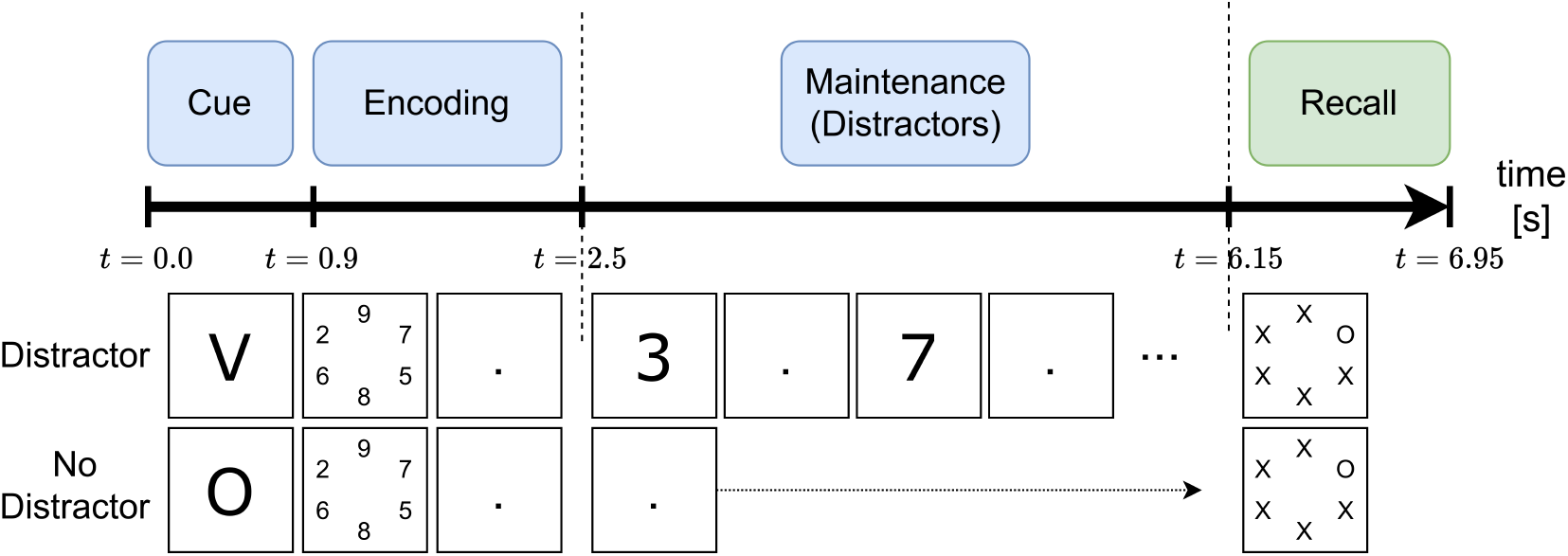
Schematic of the ‘distracter’ and ‘no distracter’ experimental trials with phases of WM. The Recall phase of interest is highlighted in green.

Brain activity was recorded using a 275-channel axial gradiometer MEG system (CTF MEG Systems, VSM MedTech Ltd) at 1,200 Hz in a magnetically shielded room. Three fiducial coils were placed at the participant’s nasion and both ear canals, which served as anatomical landmarks for offline co-registration with structural magnetic resonance imaging (MRI) scans. T1-weighted anatomical scans were acquired with a 3T MRI system (Siemens, Erlangen, Germany).

### 2.2 Problem statement and proposed approach

We consider four brain regions of interest (ROI) relevant to visual attention and WM: primary visual cortex (V1), left intraparietal sulcus (L-IPS), left and right dorsolateral prefrontal cortex, denoted by L-dlPFC and R-dlPFC. Both left and right dlPFC are thought to be involved in WM processes [30, 31], with distinct roles for the two regions [32–34], preventing combining them into a single node. The primary visual cortex V1 is a key focus of many visual attention studies, as in the original study [3]. Because visual information was presented centrally in the visual field, left and right V1 are combined into a single node. The region IPS is a candidate for the origin of top-down attention signals to V1 [35–37], and the left lobe in particular was identified in the original study as having a notable increase in alpha power just before the recall phase [3].

To capture the interaction dynamics between the four regions of interest, the state *s* is defined to represent the activity in these regions,

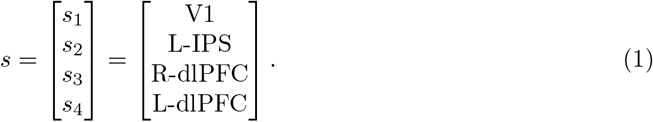

The output of the system is then directly related to these states, as the activity of these brain regions is available by the source reconstructed data for each of the brain regions, that is

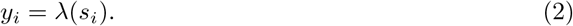

The recall period of the experiment is the focus of this study. There is no stimulus input during this phase, with the exception of the recall probe at the beginning of the period which is treated as an initial condition perturbation. To capture the internal change in dynamics in the model, the dynamics are split into a baseline model and an attention model,

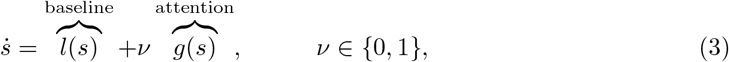

where the attention dynamics are (de)activated by the binary parameter *v*.

During the trials without distracters, the attention mechanisms are not active (*v* = 0) because no distracting input is expected. Only the baseline dynamics are active, giving,

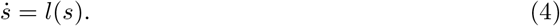

This baseline model represents the brain under baseline recall conditions, and existing literature can inform the structure of this function. During the distracter trials, the attention mechanisms are active (*v* = 1) because a distracting input is expected. This gives the dynamics

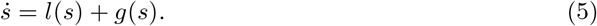

The function *g*(*s*) represents the change in dynamics which occurs when attention is internally activated. The mechanism for attention is currently unknown, so there exists significantly less literature to inform the structure of this function.

### 2.3 MEG data processing

This project uses a previously collected dataset. MEG data were preprocessed offline and analyzed using the MATLAB FieldTrip toolbox [38]. The preprocessing steps included down-sampling from 1,200 to 300 Hz, notch filtering for removal of line noise and harmonics, and independent component analysis for removal of artifacts from heart and eye movements [3]. The linearly constrained maximum variance (LCMV) beamformer approach [39] was used to obtain source reconstructed time series for the regions of interest (ROI). Volume conduction models were constructed based on single-shell models [40] of individual participant’s anatomical MRIs. Dipole positions were defined using a cortical surface-based mesh with 15,784 vertices created using Freesurfer v6.0 (RRID: SCR 001847) and HCP workbench v1.3.2 (RRID: SCR 008750). The vertices were grouped into 374 parcels based on a refined version of the Conte69 atlas [41], allowing us to reduce the dimensionality of the data, similar to [42].

Neural response time series of each anatomical parcel were computed by taking the average time series across all dipole positions within the parcel. We then retained the parcels belonging to our regions of interest (averaging over parcel time-series within each ROI), and computed power spectra for each of the ROIs using a fast Fourier transform (FFT) approach (a single subject example is given in Figure 2). Spectral analysis (5–25 Hz) focused on the *recall* window, i.e., the probe delay, which was 0.8 s in duration (giving a frequency resolution of 1.25 Hz).

**Fig 2.**
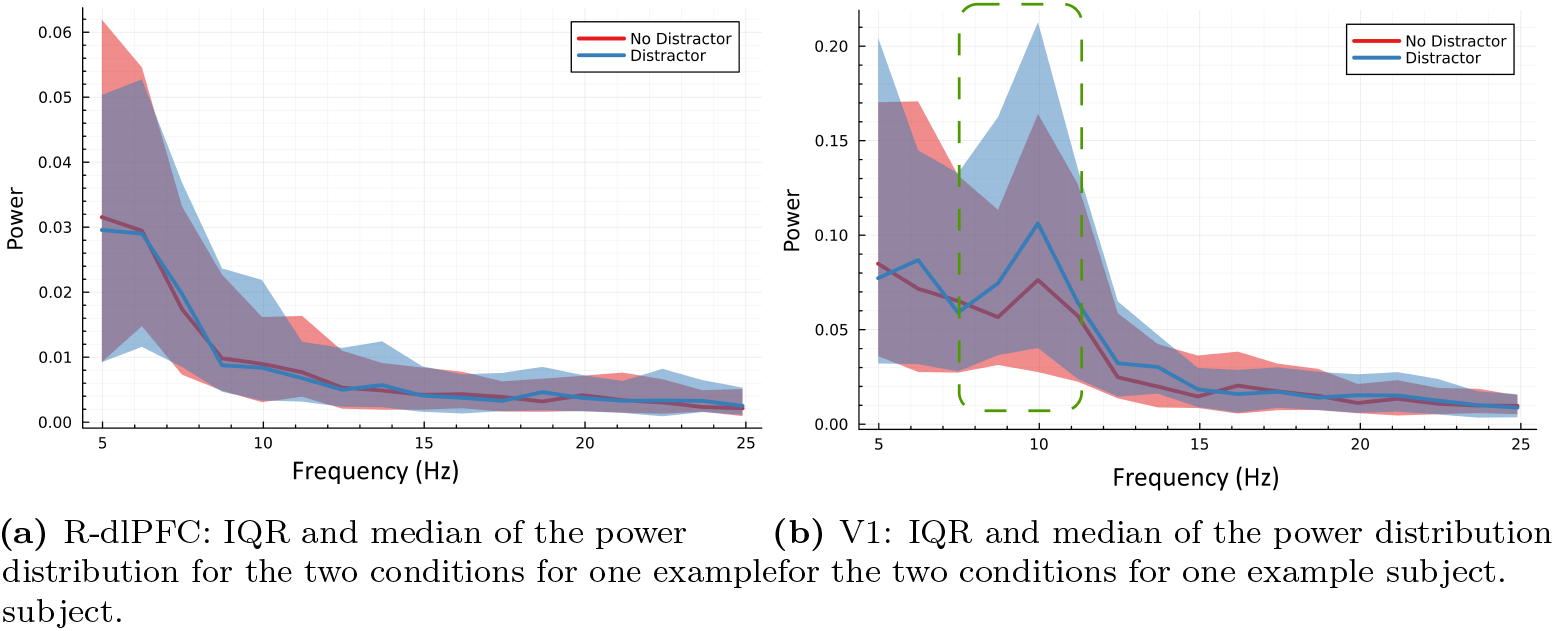
IQR (interquartile range), representing the middle 50% of the data, is denoted by the shaded regions and provides an indication of the full power distribution. Note that for V1, there is a clear increase in alpha band activity in the distracter case (circled in green), while the R-dlPFC spectral behaviour remains largely the same.

Prior to computing the FFT, a Hanning window was applied to mitigate the effects of spectral leakage. The spectra were then normalized by the maximum power over all frequencies and trial types. The median of each trial type was used as a measure of central tendency, given the skewed unimodel distribution across trials.

### 2.4 Known cortical oscillators models to capture the baseline dynamics

Many dynamical models exist to represent the activity of neuronal populations or to replicate behaviors seen in neuroimaging recordings. In a data-driven modeling context, these models trade off expressive power, i.e., flexibility to be able to represent the data well, simplicity, i.e., ease of training due to the loss landscape shape, and biological intepretability, i.e., biophysical versus phenomenological structures. Models based on populations of spiking neurons tend to be more representative of biological structures, but their highly nonlinear nature makes them difficult to fit to data without a reasonable initial guess for the parameters. On the other hand, more abstract oscillator models [43] can fit neuroimaging data well, but can be difficult to map to biological processes.

To capture this range, three candidate models are selected. The linear harmonic oscillator is one of the simplest systems that exhibits oscillatory behaviors and has been successfully used to replicate neuroimaging recordings [16]. The Duffing-Van der Pol oscillator is a combination of two well-known nonlinear oscillators, and was developed to better replicate the frequency spectra seen in EEG recordings [44]. The Wilson-Cowan model is a staple of neuronal population dynamics, reflecting known interactions between different types of neuronal populations which give rise to oscillatory behaviors [45]. Table 1 below summaries the characteristics of the chosen models.

**Table 1.** Characteristics of the three candidate baseline models.

| Oscillator Model | Expressive Power | Simplicity | Biological Interpretability |
| --- | --- | --- | --- |
| Linear Harmonic | Medium | High | Low |
| Duffing-Van der Pol | High | Medium | Low |
| Wilson-Cowan | High | Low | Medium |

### 2.5 Nonlinear function identification and symbolic regression

To reveal the attention dynamics, we use function identification by combining scientific machine learning, i.e., using neural networks as a universal function approximator, and symbolic regression. In this research, we only use feedforward neural networks, and any subsequent mention of neural networks refers to this class. The universal approximation theorems in machine learning have essentially shown that there exists a set of parameters for a neural network such that the network can approximate a wide range of functions, provided the network contains sufficient depth and width [23]. The field of scientific machine learning aims to merge the use of these universal function approximators with expert knowledge in an effort to better restrict the function identification problem [24, 25]. One development of particular interest in dynamical modeling is the neural ordinary differential equations (NODE) [26], and its subsequent universal differential equations (UDE) extension [24]. Briefly, NODE are a class of ODEs in which the dynamics of a state *x* are defined by a neural network *N*, with parameters *λ*, giving for example,

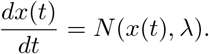

In the UDE extension of NODE framework, the differential equation is not assumed completely unknown. Instead, known differential equations are utilized along with the neural network which represents only the unknown part of the model. This has the advantage of reducing the complexity of the relationship the neural network must learn and fits our modeling approach in (5).

Ideally, the neural network function could be distilled into an arithmetic expression, which would then offer greater scientific insight into the dynamics. This simplification of the neural network to an arithmetic expression relating two datasets can be achieved through symbolic regression. In this work, we use the sparse identification of nonlinear dynamics (SINDy) method [46] composed of two steps: (1) a library, denoted by Θ(*X*), of candidate functions from which the expression for the network will be built, (2) a large set of input/output data from the network. Provided the library is rich enough, it can be assumed that the function *f* (*x*) consists of a small selection of these functions. Selection of the library is a challenging step in this process. Our choice of library is detailed in Section 3. The symbolic regression task is then to find a sparse weighted summation of the library functions, with weights denoted by Ξ, such that this sum matches the measured *Y* to within some tolerance *ϵ*, for instance by the sequential thresholding least-squares (STLSQ) method [46],

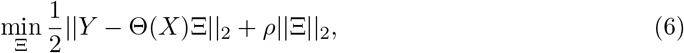

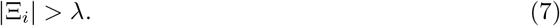

where the hyperparameters *λ* and *ρ* must be tuned to provide a reasonable balance between sparsity of the solution and error between the measurements and sparse summation.

### 2.6 Software and training algorithms

The full training algorithm is developed in the Julia programming language, primarily in the SciML framework [24] which has been used successfully in several physics-informed machine learning studies [27–29]. In particular, the DataDrivenDiffEq package is used to solve the symbolic regression problem and the SciMLSensitivity package is used for differentiation of the UDE.

## 3 Results

In this section, we first present the network structure and identified dynamics for the baseline model capturing the dynamics of interconnected ROIs corresponding to WM experiment in the absence of visual distraction. Second, we present a detailed account of our modeling and identification algorithm. We then continue validating the obtained network dynamics in both conditions, with and without distracters, by contrasting data with the model output. Finally, we identify the causal pathway, i.e., the directed neural pathway, which is obtained by comparing the models associated with each of conditions: with and without distracters.

### 3.1 Baseline oscillatory network model

To capture the baseline dynamics, we first construct three candidate networks using three types of known cortical oscillators’ models, as described in Section 2.4 and compare their performance. We compare three network models:

1. An all-to-all network composed of four, one per ROI, Wilson-Cowan oscillators
2. An all-to-all network composed of four, one per ROI, Duffing-Van der Pol oscillators
3. An all-to-all network composed of four, one per ROI, hypercolumns each composed of two harmonic oscillators.

This Duffing-Van der Pol oscillator is an extension of a nonlinear oscillator designed to replicate the wide-band power spectra of EEG signals [44]. The Wilson-Cowan oscillator is a classical neuronal population model [45, 47], which also acts as a nonlinear oscillator. The original formulation of the model includes many parameters, which makes optimization challenging. Instead, a simplified model based on the deterministic part of the model used in [48], is used here. Detailed mathematical models for the Duffing-Van der Pol and Wilson-Cowan models are provided in Section 5.

The third network is our design inspired by the columnar models of the cortex [49, 50], where we replaced the abstract minicolumn model with a harmonic oscillator. We refer to this network by *Double Harmonic Oscillator* in the rest of the article. The coupling architecture is shown in Figure 3. Each linear oscillator in the region is fully connected to corresponding, i.e., low or high frequency, linear oscillators in the other regions, forming two fully connected networks. The training algorithm is designed such that one of the networks captures low-frequency behaviors and the other captures high-frequency behaviors. There also exists cross-coupling between the networks, but not within the regions. This extension preserves the linearity of the model, while increasing its expressive power by increasing the number of autonomous oscillators in the network and allowing for more complex connectivity.

**Fig 3.**
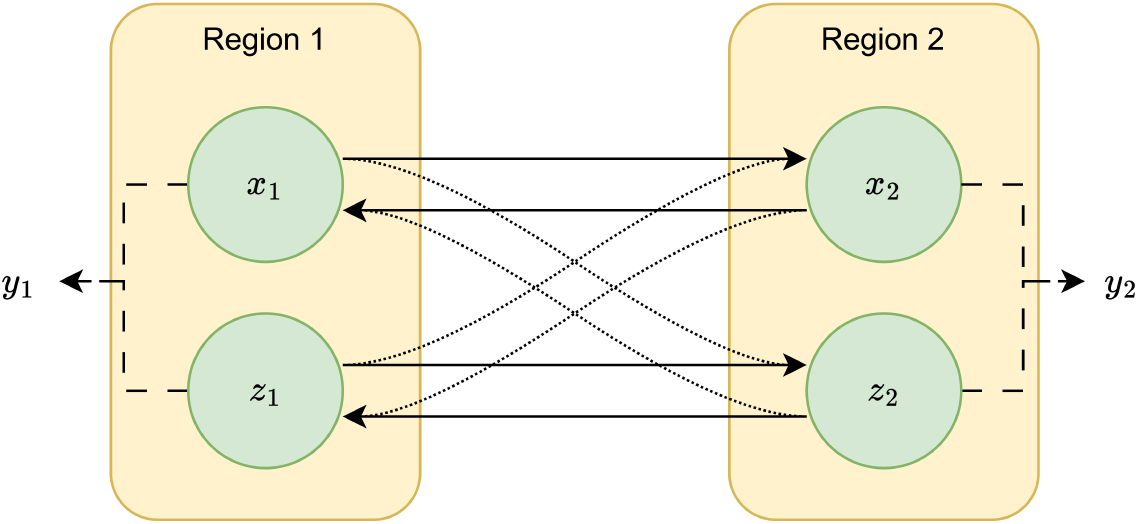
A two-region schematic of the connectivity of the double harmonic oscillator network. The solid black lines denote couplings for the two fully connected networks. There also exists cross couplings between the networks, denoted by the fine dotted lines.

Each region, i.e., ROI, *i* ∈ {1, …, 4}, consists of two oscillators, represented by states *x*_*i*_ and *z*_*i*_. Linear diffusive coupling within the networks *x* and *z* have strengths *k* and *c*, respectively. Influence of the *z* network on the *x* network occurs via linear diffusive coupling of strength *b*, and vice-versa with strength *l*. This gives the dynamical model,

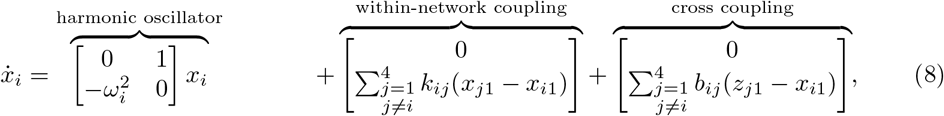

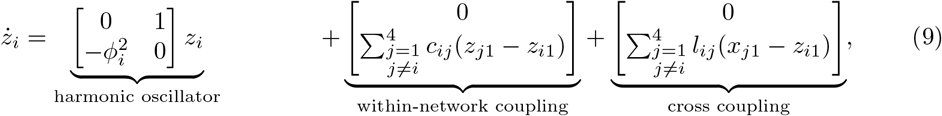

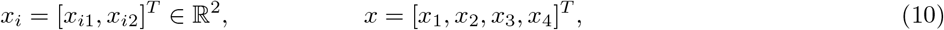

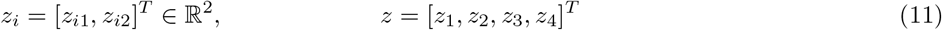

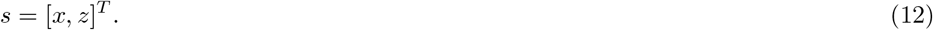

The model outputs are the sum of the first states of the two oscillators in each region,

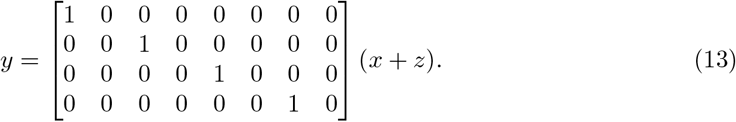

We compare the performance of the three network models mentioned above to choose the baseline model. The manner of comparison is explained in Section 3.2.1. The coupled linear oscillator network developed in this study achieved the best performance as shown in Table 2.

**Table 2.**
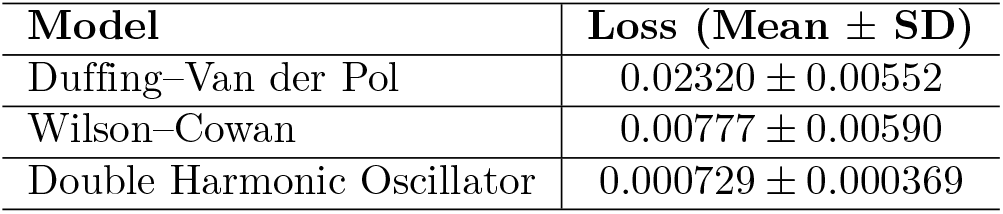
Baseline model performance comparison. Mean and standard deviation of the loss function value (39) with optimal parameters, taken across the four participant models.

| Model | Loss (Mean ± SD) |
| --- | --- |
| Duffing–Van der Pol | $0.02320 \pm 0.00552$ |
| Wilson–Cowan | $0.00777 \pm 0.00590$ |
| Double Harmonic Oscillator | $0.000729 \pm 0.000369$ |

### 3.2 The Modelling and Identification Algorithm

In this section, we first provide an overview of our oscillatory network modeling algorithm. Then, each step of the algorithm are detailed and explained.

#### Algorithm 1

Data-driven oscillatory network modeling with condition-dependent coupling laws together with causal pathway identification method

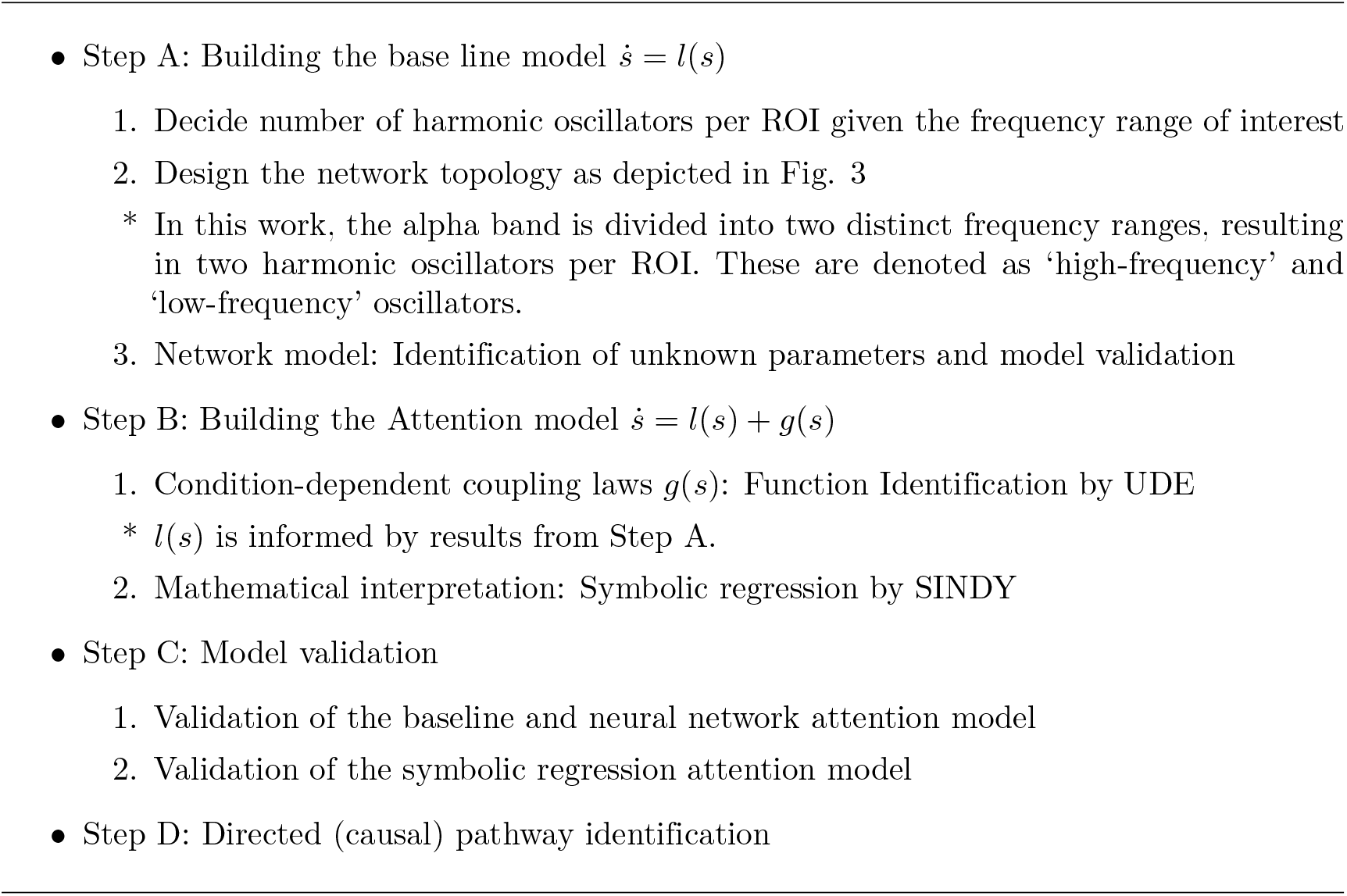

#### 3.2.1 Step A: Building the Baseline model

As discussed earlier, we found that the high-dimensional linear oscillator best represents the network model *ṡ* = *l*(*s*) for the condition without distracters. In this work, we focus on the alpha band (5 to 25 Hz) and adopt a double harmonic oscillator model for each region, dividing the frequency band into two harmonics corresponding to 10 Hz intervals. A single harmonic oscillator only adequately captured data within the 5 to 12 Hz range, prompting us to utilize two harmonics. While it’s possible to add more harmonics, the choice should balance the complexity of parameter fitting with accuracy in data representation.

The large number of parameters and states, especially in the double harmonic oscillator case, make the problem prohibitive to solve directly. Instead, a multi-stage training algorithm is developed to narrow down the parameter search space. Generally, parameter fitting proceeds with a multi-level single linkage (MSLS) algorithm [51] with local SLSQP, followed by Adam optimization [52], allowing for efficient global search of the complex but continuous and constrained parameter search space. This process also holds for all oscillators we tested in Table 2. For the double-harmonic oscillator, the process is detailed as follows. First, the high- and low-frequency networks are independently fit to the high- and low-frequency portions of the power spectrum. To fit the parameters of the full network, including both networks and cross-coupling, the results from the previous optimization results are used as initial conditions, with cross-coupling initialized at zero. This allows a local optimization algorithm, here Adam, to be used, as the parameters are already nearby a reasonable solution. Complementary information are provided in Section 5.

#### 3.2.2 Step B: Attention Model Identification

With selection of the baseline model complete, we move to the attention model identification, which captures both the dynamics, i.e., with and without distracters. The full optimization problem can be found in the supplementary information (45). The key difficulty in solving this problem is the large number of parameters introduced by the embedded neural network model, but the problem is made tractable through use of the Gauss adjoint sensitivity method [24] and a warm start of the baseline model parameters from the baseline model fitting.

The result of the optimization is a baseline model *l*^*\**^(*s*) which captures the nominal WM recall dynamics, and a neural network *N*^*\**^(*s*) which captures the change in these dynamics due to activation of attention mechanisms.

To determine what neural mechanisms the function *N*^*\**^(*s*) is describing, the symbolic regression technique SINDy, is used to construct an arithmetic expression which captures the input-output behavior of *N*^*\**^(*s*). The first step in this process is the creation of the input-output dataset. This is generated by using the trajectory of the distracter (*v* = 1) condition as the input set, and passing these points through the network function,

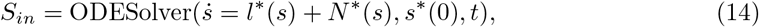

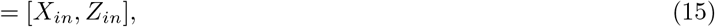

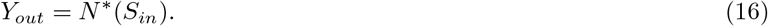

This selection of the input set guarantees that the full input-output set captures the most important relationship learned by the neural network, as this is the trajectory that network is trained on.

The next step is the selection of the library of candidate functions. This library includes a monomial expansion of the network inputs from degree two to degree five, plus a collection of diffusive coupling terms as is used in the baseline model. The network function *N*^*\**^ is a collection of eight separate networks, each with a different set of inputs. This structure means that the library of candidate functions is also different for each of the separate networks, and that the sparse optimization must be completed separately for each network. As an example, the library for the network 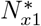 is

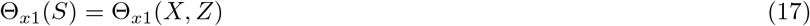

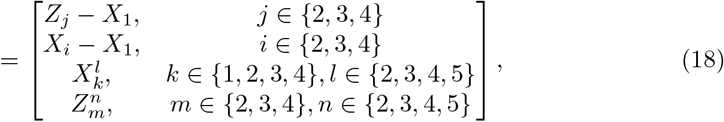

including transformations of the same inputs as the function *N*_*x*1_. The sparse identification problem is then

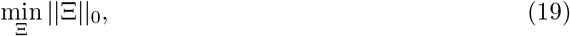

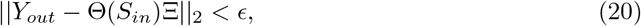

which is solved using the STLSQ algorithm. The result of solving all eight sparse identification problems is a new function which is roughly equivalent to the network function but with significantly fewer terms and parameters,

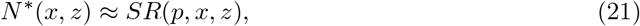

where *p* are the nonzero elements of the sparse vector Ξ^*\**^. To ensure optimality of the parameters, the optimization problem for the full dataset defined in (45) is repeated with this model structure, for detailed information see Section 5,

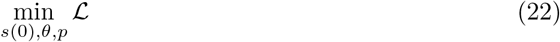

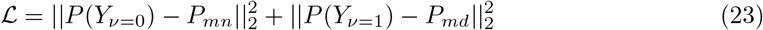

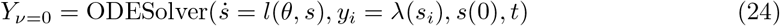

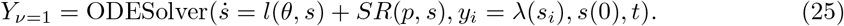

#### 3.2.3 Step C: Model validation

The described method is used to fit four independent models on four separate subject datasets. Here, the model outputs are compared to the observed data.

### Validation of the baseline and neural network attention model

A comparison of the data and model output for one subject V1 is shown in Figure 4. Visual inspection of the fit demonstrates that the baseline model captures the data in this condition well. Together, the baseline plus the neural network also capture the data corresponding to the condition with distracter well, suggesting that the network can replicate the attention dynamics. Similar agreement is found for the other three subjects and regions.

**Fig 4.**
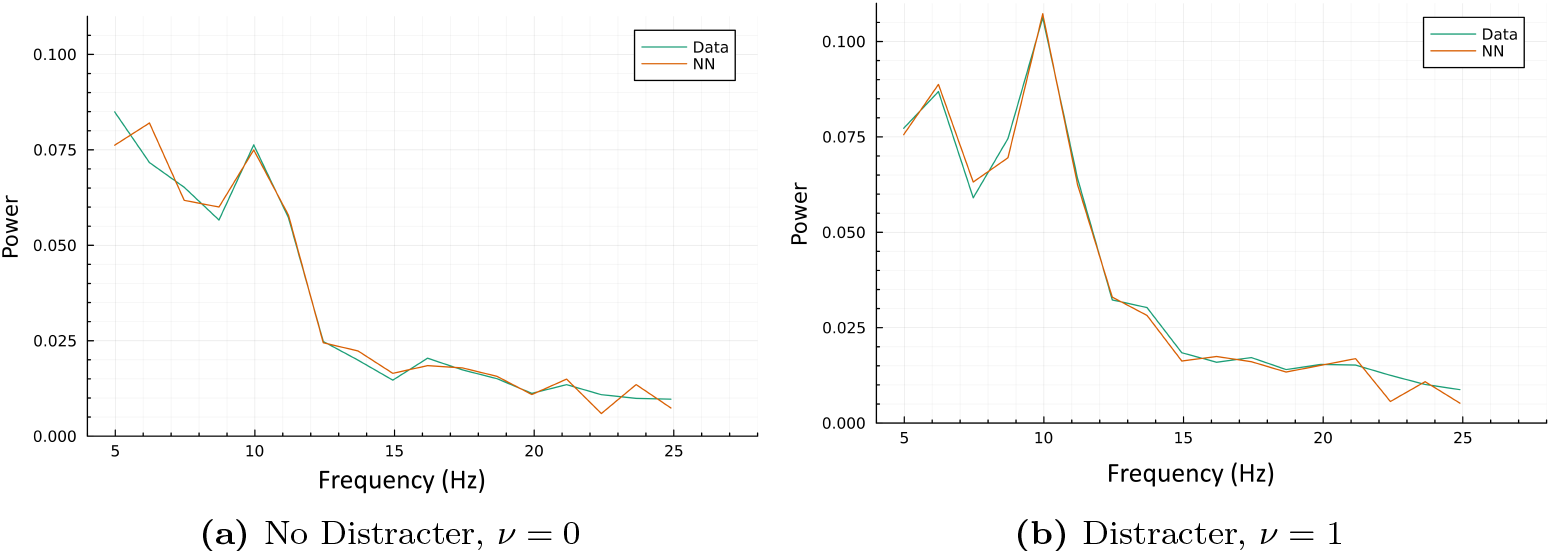
Fit of the UDE, with baseline double extended linear oscillator and attention neural network models, to the observed data for Region V1.

### Validation of the symbolic regression attention model

A comparison of the data, neural network model, and symbolic regression model for all four regions of one of the subjects is shown in Figure 5. The close agreement between the neural network and symbolic regression models demonstrates that the symbolic regression technique is able to reduce the highly parameterized network into a simpler expression which captures the same behaviors. Similar agreement is found for the other three subjects.

**Fig 5.**
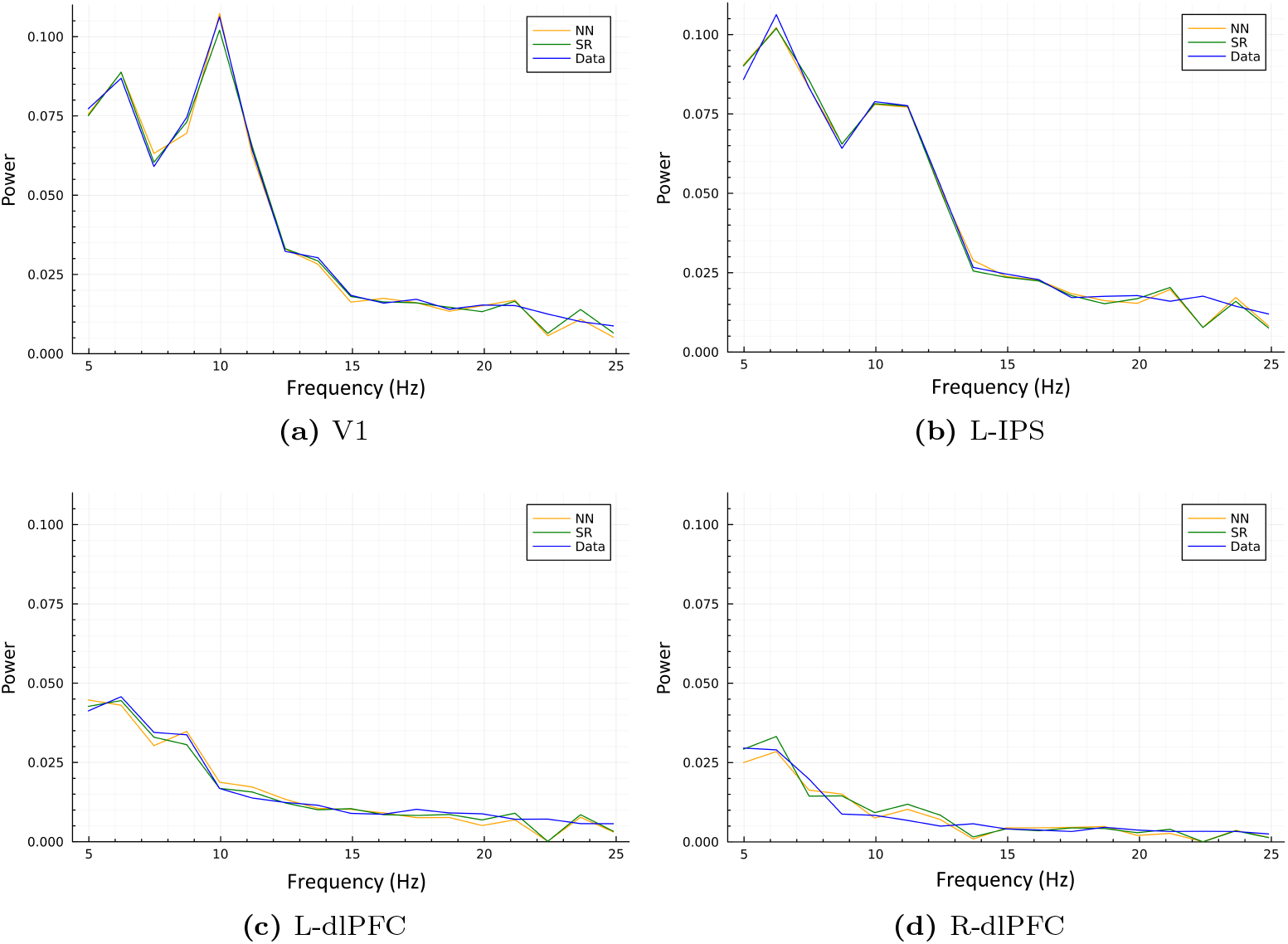
Comparison between the symbolic regression and neural network models in capturing the observed data corresponding to condition with distracters.

### 3.3 Step D: Directed pathway identification

This section presents our method to infer directed pathways assumed to be activated in the presence of distraters. The distilled attention-activated dynamics for a single participant are provided in. A reminder that the state is defined as

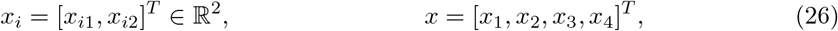

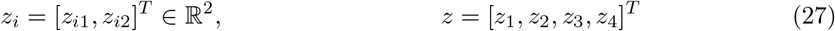

where *x* are the low-frequency oscillators and *z* are the high-frequency oscillators. The ROIs V1, L-IPS, L-dlPFC, and R-dlPFC, are numbered by 1 to 4, respectively.

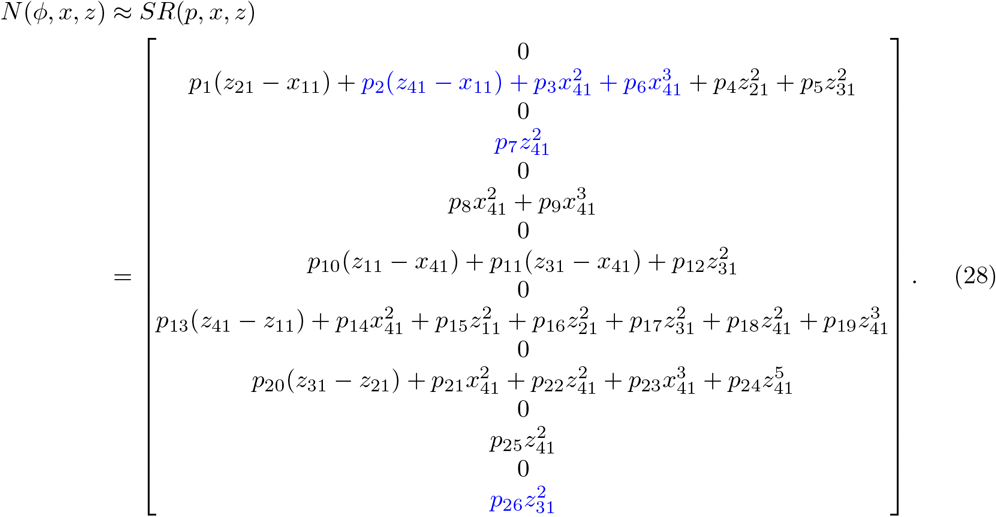

To aid interpretation of the symbolic regression models, Figure 6 provides the value of the non-zero terms of *SR*(*x*(*t*), *z*(*t*)) evaluated over the solutions *x*(*t*), *z*(*t*) of the full model for one of the subjects. These terms indicate the impact of the attention mechanism, here represented by the function *SR*(*x, z*), on the dynamics of the oscillators over the course of the solution trajectory. In particular, visualization of these terms allows for an assessment of which states in the model are most impacted by the activation of the attention model. From this, it is clear that Region V1 experiences significant changes in dynamics with the activation of the attention mechanisms, as both the high- and low-frequency *SR* function terms for this region are large in magnitude over the course of the solution trajectory. A similar trend holds for R-dlPFC. The condition-dependent coupling terms in the dynamics of V1 and R-dlPFC are highlighted in blue. We see that V1 dynamics is influenced by the sstates of region 4, that is R-dlPHC. Moreover, the new term in the dynamics of R-dlPFC is the state of the L-dlPFC. Figure 8 shows the directed, i.e., causal, distracter-dependent pathway for this subject.

**Fig 6.**
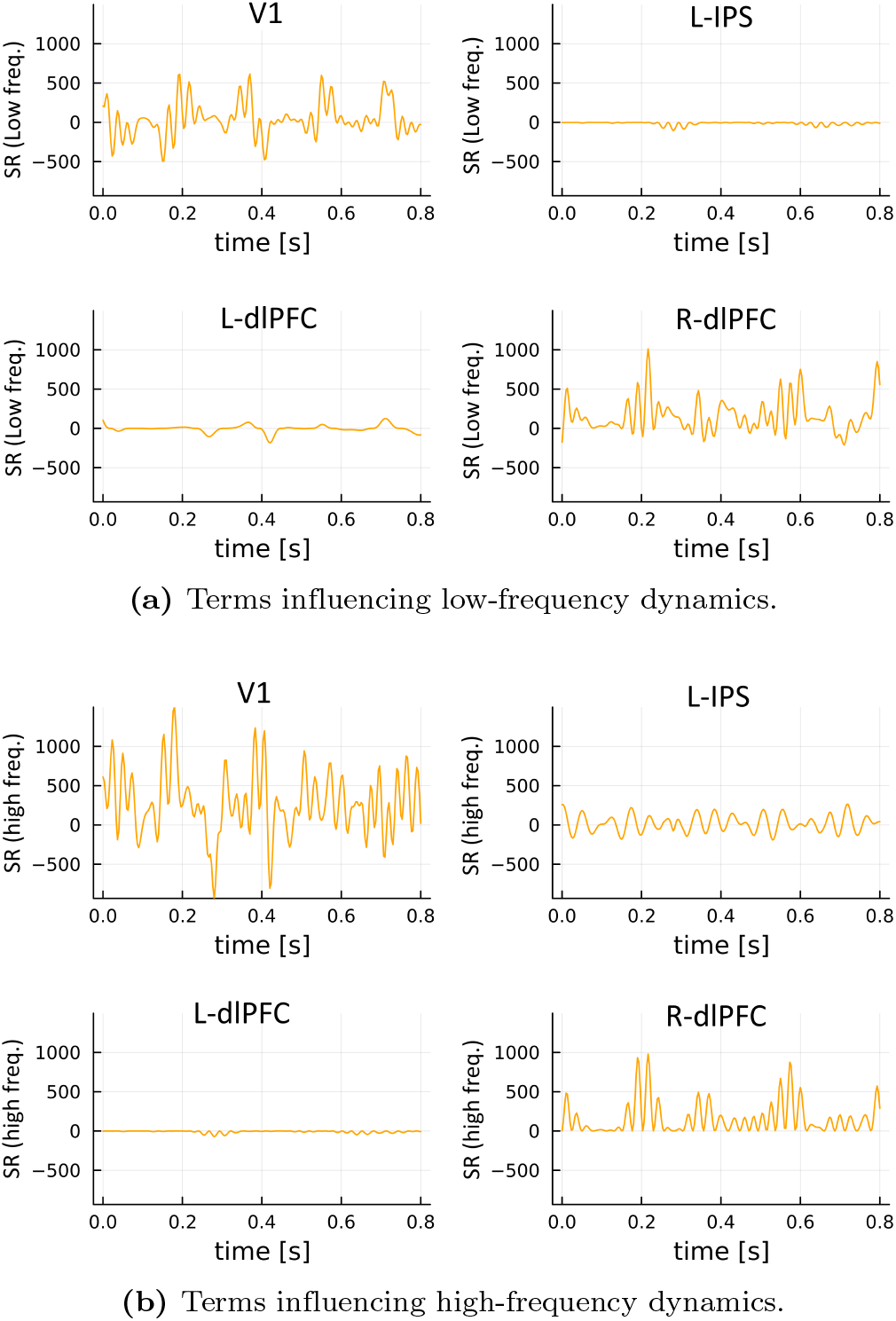
*SR*(*x*(*t*), *z*(*t*)) evaluated over the solutions *x*(*t*), *z*(*t*) of the full model.

**Fig 7.**
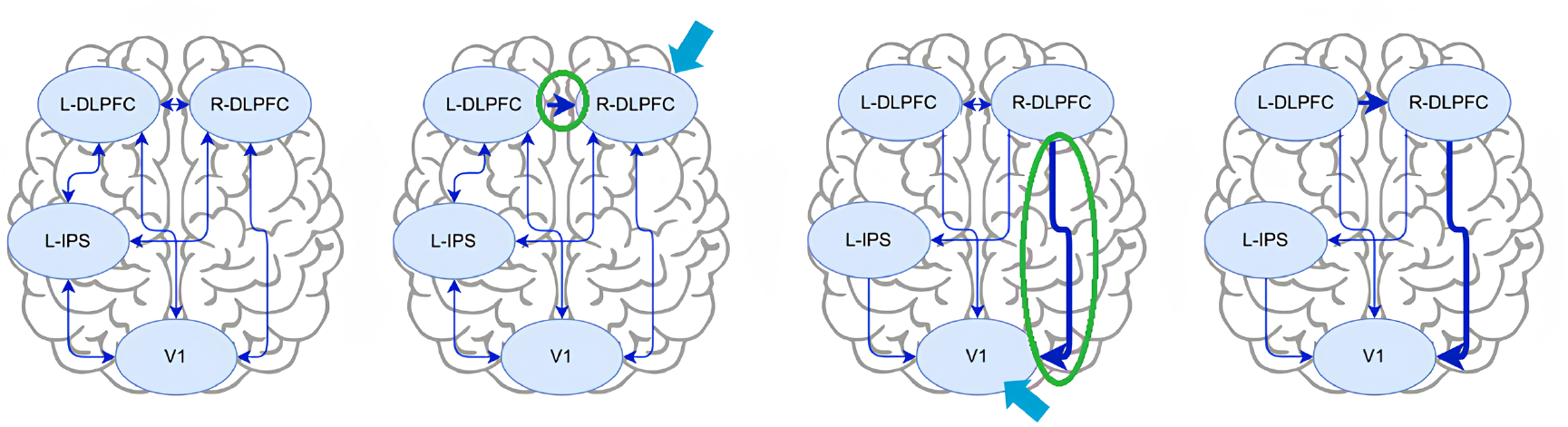
Identifying the cause in changes of the dynamics of the two ROIs (V1, and R-dlPFC) which experienced major changes as shown in Fig. 6

**Fig 8.**
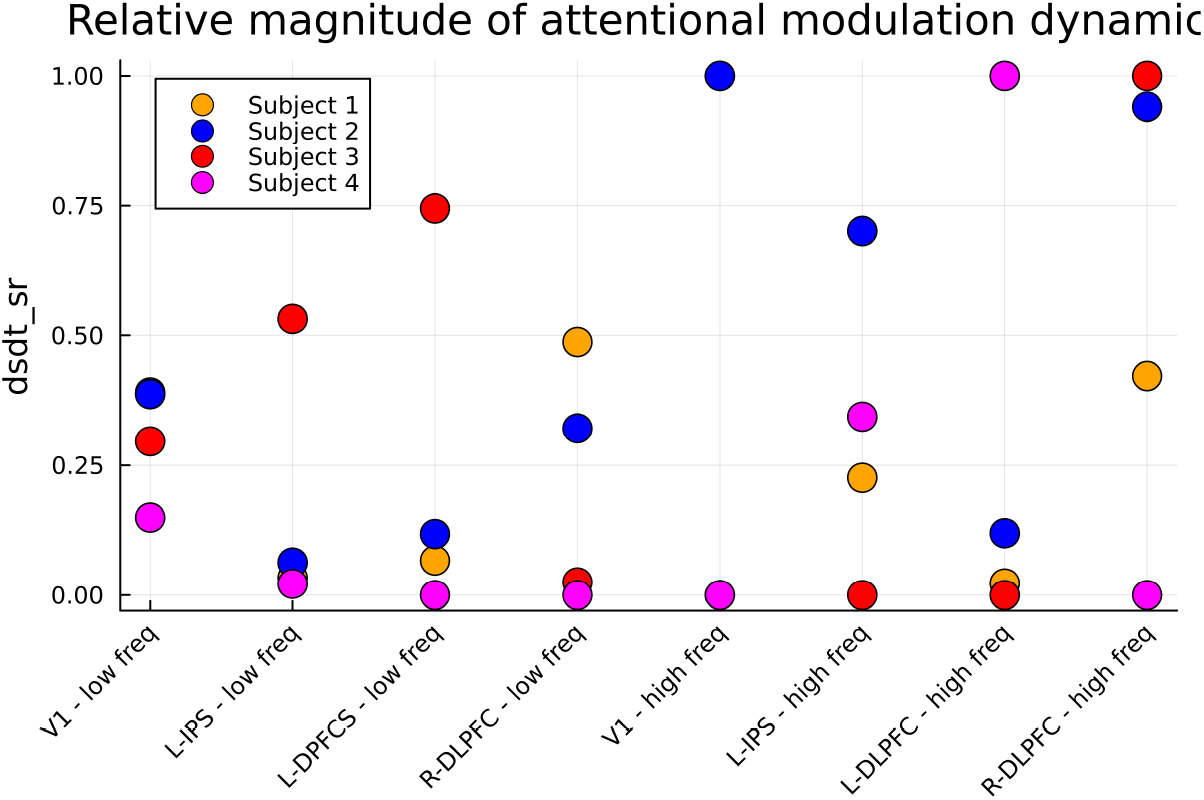
Relative magnitude of the attention modulation dynamics (|*SR*(*x, z*)|) over all regions, participants, and frequency ranges. High magnitudes indicate that the dynamics of a region are heavily impacted by the activation of attention mechanisms.

### 3.4 Distrater-dependent causal pathway

In this section, we analyze the four independent subject models, which together suggest the emergence of a distracter-dependent pathway from the dlPFC to V1. Neural mechanisms are identified through examination of the optimized symbolic regression attention models. The results of all four subjects corresponding to the ROIs with major changes due to distracters are provided in the Supplementary information. Although there is subject-to-subject variation, we observe that there exists a consistent moderate increase in the dynamics of low frequency V1 when attention mechanisms are activated. This aligns well with existing literature on visual attention [3, 53–55]. To determine what influences cause this large change in the dynamics of low-frequency V1, denoted by state *x*_1_, on activation of attention mechanisms, the symbolic regression expressions for this region for all four participants, *SR*^1^ to *SR*^4^, are presented below

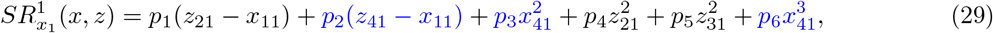

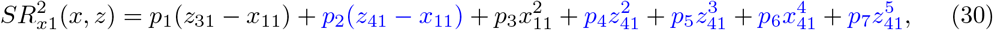

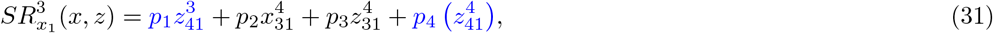

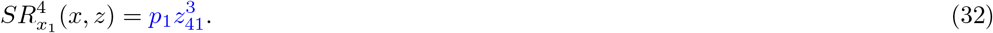

Terms colored in blue correspond to R-dlPFC, note their prominence in all four participant expressions. This suggests that input from the R-dlPFC is important to the change in behavior of V1 when distracters are anticipated. This observation is especially interesting considering that power spectral analysis of R-dlPFC reveals minimal changes between conditions with and without distracters (see comparison between R-dlPFC and V1 in Figure 2). Thus, although data analysis suggests that the R-dlPFC is not active in attention mechanisms, this dynamical analysis suggests that the region plays an important role in cognitive control of other regions.

## 4 Discussion

In this section, we first highlight the novelty of our contributions and provide a comparison with the existing approaches, and then discuss its limitations and future perspectives.

## 4.1 Novelty and Comparison

We have compared several baseline models for their ability to replicate the power spectra of measured MEG signals within a data-driven modeling framework. Our results show that a higher-dimensional network of linear oscillators, one per brain region, outperforms a lower-dimensional network of nonlinear oscillators for this task, yielding a lower fitting error. This finding highlights the importance of balancing model complexity with careful formulation of the optimization problem in data-driven modeling applications. More specifically, model selection should be guided by both physical understanding of the system and simplification of the loss function. In this work, we have introduced a baseline high-dimensional linear oscillator network model that mimics a cortical hyper-column composed of a small number of harmonic oscillators, here, two oscillators, enabling the representation of a broader frequency range. Additionally, we have introduced a multi-stage parameter-fitting procedure that enables the network to capture broad frequency behavior, a critical aspect of MEG modeling studies.

Furthermore, We have extended previous work in the scientific machine learning domain by applying it to MEG data. Our results demonstrate that neural networks, in combination with the baseline model, can capture changes in the system’s response, validating the existing universal differential equations (UDE) framework. Furthermore, symbolic regression techniques, specifically SINDy, previously used in the literature alongside UDEs, were shown to accurately replicate the behavior learned by neural networks, supporting their use as tools to improve the interpretability of neural network models. However, the dynamic interpretation of the neural network’s mathematical expression via symbolic regression was not evident. Therefore, we developed a novel method to interpret it by measuring inter-regional influences expressed by monomial functions.

We have aimed at providing a directed pathway identification framework in which two conditions, here with and without distracters, can be compared in terms of their underlying mechanisms. Our proposed data-driven modeling and identification algorithm allows causal pathway detection through identifying nonlinear coupling functions. Omitting the latter stage, that is, fitting the two conditions, with and without distracters, to the baseline oscillatory model, would give two models with the same model structure, but different coupling weights. We also verified this approach and found that, per participant, comparing weights between these two models (see Supplementary information) does not yield a clear distinction in inter-regional coupling between the two conditions.

The above comparison demonstrates that although both models replicate the data well, the proposed algorithm’s flexibility enables data-driven discovery of models that suggest previously unrecognized mechanisms underlying attention dynamics. For example, our algorithm revealed an L-dlPFC*→* R-dlPFC*→*V1 pathway that may play an important role in attention dynamics, yet this pathway is entirely absent in the hypothesis-driven model which relies on comparing coupling weights. This insight can serve as a starting point for further investigation in future empirical studies.

## 4.2 Limitations and perspectives

Despite inter-subject variability, data from all four participants examined here showed the emergence of a distracter-dependent pathway from dlPFC to V1. This finding is consistent with the established role of the dlPFC in cognitive control and suggests that distracter processing recruits a directed interaction from prefrontal to visual regions. These preliminary findings demonstrate how our modeling approach can be used for the generation of hypotheses which can then be tested in subsequent empirical studies.

A limitation of the current framework is that inter-subject variability is not explicitly handled in the model. Instead, a separate model is fit to each subject, restricting coverage of the network input domain and increasing the risk of overfitting, in addition to increasing complexity of subsequent model analysis. Extending the algorithm to multi-subject datasets would improve training robustness by forcing the network to capture dynamics that are consistent across subjects, thereby enabling more generalizable insights into condition-dependent tasks, such as the attention-working memory interplay.

The proposed algorithm currently focuses on reproducing static frequency-domain characteristics of MEG activity. Although these features are relevant to attentional dynamics during recall, transient time-domain properties are also likely to play an important role and could be incorporated through refinement of the optimization metrics used during training. In addition, neuroscientific studies commonly analyze behavioral measures, such as reaction time and response accuracy, alongside MEG recordings to obtain a more comprehensive understanding of cognitive processes. These behavioral variables are not included in the current framework, but could be integrated through additional, potentially continuous, activation variables or by extending the model outputs. Such developments would further strengthen the connection between experimental neuroscience and dynamical systems modeling.

## 4.3 Conclusion

This study introduced a phenomenological data-driven framework for oscillatory network modeling that learns condition-dependent coupling laws directly from MEG recordings. Additionally, an analysis framework was presented to identify the directed neural interactions underlying changes in the dynamics of working-memory recall in the presence of distracters. The algorithm provided a systematic approach for deriving data-driven insights into neural mechanisms in settings where strong mechanistic hypotheses were unavailable. First, a hyper-column linear oscillatory network model was developed to capture interactions between brain regions of interest in the condition without distracters. Then, universal differential equations coupled with symbolic regression were used to identify changes in recall dynamics under the condition with distracters and present them in a biologically interpretable form.

Application of the algorithm to four participants revealed a consistent pathway from dlPFC to V1, supporting the dlPFC’s role in regulating visual recall and distracter processing. In general, the study highlighted the potential of data-driven dynamical systems approaches, especially function-identification techniques such as UDEs, for generating hypotheses in cognitive neuroscience, while also identifying several directions for developing dynamical systems models suited to experimental neuroscience.

## 5 Supplementary information

In this section, we provide more detailed descriptions of the models, algorithms and results found in this study.

### 5.1 White-box models of interconnected cortical oscillators

Here we provide detailed mathematical models of the three white box models explored to select the best fit for the base line cortical network composed of four regions. Each of the four regions is represented by an oscillator. The oscillators are assumed to be communicating over a fully connected, all-to-all, graph.

#### Duffing-Van der Pol Oscillator

This Duffing-Van der Pol oscillator is an extension of a nonlinear oscillator designed to replicate the wide-band power spectra of EEG signals [44].

Each region *i* ∈{1, …, 4} contains a self-sustaining Van der Pol oscillator with unique parameters *l, µ*, which together determine the uncoupled frequency of the regions. The network is fully connected via Duffing oscillator coupling. This coupling includes a diffusive linear term with strength *k* plus a diffusive cubic term with strength *b*. This gives the dynamical model,

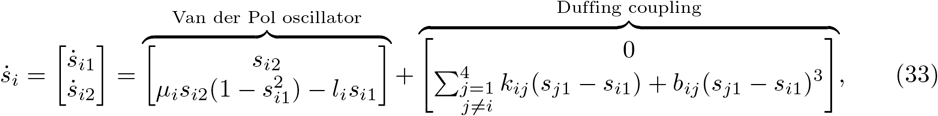

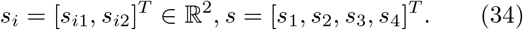

The model outputs are the first state of each oscillator. When fitting the model, its predicted power was consistently lower than the data. To account for this, a single scaling parameter was introduced. This does not affect the model’s validity because the data were normalized to their maximum power, making the absolute magnitude arbitrary. The relative power across regions and frequencies, which is the quantity of interest, is preserved.

The model has eight Van der Pol parameters and 24 coupling parameters, giving a total of 32 parameters. In addition to this, there are eight states requiring eight initial conditions, plus the power scaling parameter, meaning the full optimization problem has 41 decision variables.

#### Wilson-Cowan Oscillator

The Wilson-Cowan oscillator is a classical neuronal population model [45, 47], which also acts as a nonlinear oscillator. The original formulation of the model includes many parameters, making optimization challenging. Instead, a simplified model based on the deterministic part of the model used in [48], is utilized here. Each region *i* ∈ {1, …, 4} includes an excitatory population, represented by *s*_*i*1_, and an inhibitory population, represented by *s*_*i*2_. The oscillation of each region emerges from the interaction between the two populations. As the terms excitatory and inhibitory suggest, excitatory populations can only excite or increase the activity of connected populations, while inhibitory populations can only inhibit or decrease the activity. The inhibitory populations connect only to themselves and to the excitatory population within the same region; this occurs through a non-diffusive linear term with strength *b*. The excitatory populations also connect to the local inhibitory populations through a non-diffusive linear term with strength *l*. These external inputs are summed and pass through a logistic sigmoid function *S*. To form the greater network, the excitatory populations are fully connected through a non-diffusive linear term with strength *k*. This gives the dynamical model,

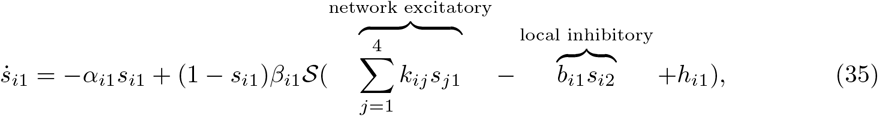

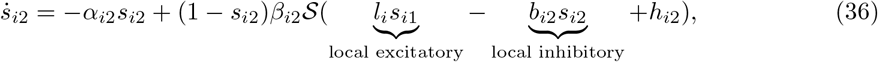

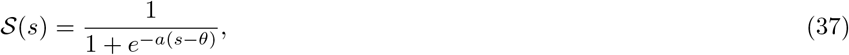

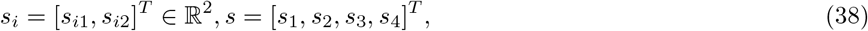

where *α*_*i*_, *β*_*i*_, *k*_*i,j*_ are positive constant parameters denoting the coefficients and coupling weights, respectively. Each population also receives a constant input *h*_*i*_, representing background input from unmodeled regions. The parameters of the sigmoid function are set at *a* = 1, *θ* = 4. MEG records the activity of excitatory populations, so the model outputs are the first state of each region. As in the Duffing-Van der Pol oscillator, a scalar parameter for the power is introduced.

This model has 16 internal dynamics parameters and 36 coupling parameters, giving a total of 52 parameters. In addition to this, there are eight states requiring eight initial conditions, plus the power scaling parameter, meaning the full optimization problem has 61 decision variables.

### 5.2 Optimization Problem Formulation

#### 5.2.1 Baseline Model

For selection of the baseline model *ṡ* = *l*(*s*) from the set of candidate functions, only the data corresponding to the condition without distracters is used. Denote the median power spectrum calculated from the data of the condition without distracters *P*_*mn*_. These are the data characteristics the model must capture. Denote *Y*_*v*=0_ as the time-domain output of the model under attention inactive conditions. These outputs are produced from a numerical ODE solver, giving the optimization problem,

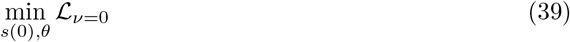

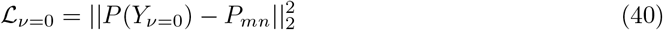

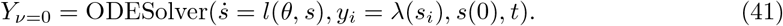

#### 5.2.2 Full Model with Attention Dynamics

The full model must replicate both the conditions with and without distracters. For the condition with distracters, the parameters of the baseline model *θ* and the parameters of the neural network *ϕ*, with the complete loss function,

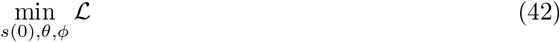

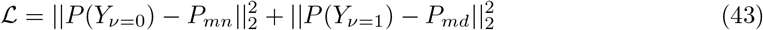

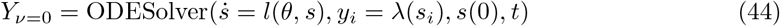

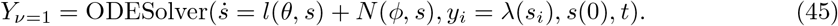

The complete optimization model is optimized using the baseline fitted parameters as an initial condition, ensuring that the no-distracter condition is well represented. The neural network parameters are initialized near zero. Because the parameters are initialized well, the optimization algorithm Adam is used to determine the optimal parameters.

The Adam optimizer requires gradients of the loss function with respect to the model parameters. Since this UDE contains a neural network with 5000 parameters, these gradients are computed efficiently using the Gauss adjoint method, which improves computational and memory efficiency over autodifferentiation and the backsolve adjoint method [24].

#### 5.2.3 Changes in dynamics of each of ROIs with distracters for all subjects

Figure 8 summarizes these results for all four participant models, giving the mean relative magnitude of these *SR* function terms for the four regions of interest in the two frequency bands.

## Acknowledgments

The authors would like to thank Luca Laurenti and Frederik Mathiesen for helpful discussions.

